# The effect of sequence mismatches on binding affinity and endonuclease activity are decoupled throughout the Cas9 binding site

**DOI:** 10.1101/176255

**Authors:** Liyang Zhang, H. Tomas Rube, Harmen J. Bussemaker, Miles A. Pufall

## Abstract

The CRISPR-Cas9 system is a powerful genomic tool. Although targeted to complementary genomic sequences by a guide RNA (gRNA), Cas9 tolerates gRNA:DNA mismatches and cleaves off-target sites. How mismatches quantitatively affect Cas9 binding and cutting is not understood. Using *SelexGLM* to construct a comprehensive model for DNA-binding specificity, we observed that 13-bp of complementarity in the PAM-proximal DNA contributes to affinity. We then adapted *Spec-seq* and developed *SEAM-seq* to systematically compare the impact of gRNA:DNA mismatches on affinity and endonuclease activity, respectively. Though most often coupled, these simple and accessible experiments identified sometimes opposing effects for mismatches on DNA-binding and cutting. In the PAM-distal region mismatches decreased activity but not affinity, whereas in the PAM-proximal region some reduced-affinity mismatches enhanced activity. This mismatch-activation was particularly evident where the gRNA:DNA duplex bends. We developed integrative models from these measurements that estimate catalytic efficiency and can be used to predict off-target cleavage.

## INTRODUCTION

The CRISPR-Cas9 system is driving the next wave of genetic research. The flexibility and relative ease of using a programmable guide RNA (gRNA) to target Cas9 to specific complementary DNA sequences has made genomic editing readily accessible (1, 2). Further, catalytically deactivated Cas9 (dCas9) has been adapted to manipulate gene expression and chromatin state, visualize chromosomal loci, and other applications (3-6). Despite active research in the area, developing rules that predict the effect of mismatches on off-target binding and endonuclease activity have had limited success, impeding broader use of Cas9.

The Cas9 ribonucleoprotein (Cas9-RNP), composed of the protein and annealed CRISPR RNA (crRNA) and trans-acting crRNAs (tracrRNA), cuts DNA in a two-step process. First the Cas9-RNP binds specific DNA sequences called protospacers (7). The Cas9 protein itself binds the sequence NGG called protospacer adjacent motif (PAM), and partially melts the DNA duplex. The gRNA, within the crRNA, then basepairs with the 20-nt protospacer DNA. The resulting RNA:DNA duplex form an R-loop that then basepairs in a sequential fashion from the PAM toward the distal region of the protospacer (8, 9). The resulting “zipping model” for target identification and binding has shown that mismatches between the gRNA and protospacer affect the kinetics and affinity of binding by interrupting zipping, with adjacent mismatches having a more than additive effect. A structural change in the Cas9 protein accompanies basepairing in the PAM-distal region, activating the endonuclease, and cutting both strands of DNA upstream of position 3 in the protospacer (7).

One consequence of the zipping model is that the gRNA:protospacer duplex can be divided into two functional domains. Basepairing within the seed, or PAM-proximal region (positions 1-10), contributes most strongly to targeting the Cas9-RNP to specific sequences. Base-pairing in the PAM-distal region (positions 11-20) does not contribute as strongly to affinity, but is important for enzymatic activity by inducing the conformational change that activates Cas9 protein (10). Thus, the current model is that the seed sequence targets Cas9-RNP to binding sequences whereas the PAM-distal region is important for the Cas9-RNP endonuclease activity (10, 11). However, the boundaries of each domain are not clear, nor are the quantitative effects of mismatches on binding, endonuclease activity, and overall enzyme efficiency within each of these regions.

The DNA-binding specificity of Cas9-RNPs has primarily been inferred from genomic localization in cells using chromatin immunoprecipitation followed by deep sequencing (*ChIP-seq*). In addition to the intended target, the Cas9-RNP binds from a handful to thousands of other sites in the genome (12, 13). These off-targets sites still retain elements of the PAM and matching protospacer, with a consensus sequence ranging from 5 to 13 bp of complementarity. Further, although generally mismatches within the PAM-distal region (positions 11-20) are better tolerated than mismatches within the seed region, the Cas9-RNP can still bind sites with some seed region mismatches. More recently, high-content kinetic studies of Cas9-RNP binding *in vitro* have shown that mismatches in the PAM-proximal region dissociate more readily than distal mismatches, but that distal mismatches can accentuate the effect of proximal mismatches (14). Although these studies are consistent, they do not present a comprehensive model for the DNA-binding specificity of the Cas9-RNP.

In addition to off-target binding, off-target cleavage by Cas9-RNPs can be prevalent (15). Measuring the prevalence of *in vivo* off-target cleavage has been challenging. Detection of cleavage in cells relies on detection of the indels, mutations caused by non-homologous end joining (12, 15-17), or by tagging double-strand breaks to mark off-target sites (18, 19). Although improving in sensitivity, these techniques still rely on cellular repair mechanisms, which vary from cell to cell. Nonetheless, it is clear that mismatches throughout the PAM and full 20-bp protospacer affect cleavage efficiency. A comparison of activities in the same cells shows a remarkable lack of overlap between the most highly bound and cleaved sites (12). Further, recent reports posit that sites with little complementarity can be cleaved by the Cas9-RNP *in vivo*, and that off-target editing is in fact widespread (18). These findings suggest a distinction between binding and cleavage that is not fully appreciated.

*In vitro* measurements of off-target cleavage have confirmed that efficient cleavage requires both an intact NGG within the PAM and complementarity throughout the protospacer. Using randomized libraries or isolated genomic DNA, thousands of off-target cleavage sites are detectable (19). Analogous to their effect on binding, mismatches appear to have a more pronounced impact on cleavage efficiency the closer they are to the PAM, and some are more deleterious than others (2, 16, 20-22). Although this reinforces the model that endonuclease activity is governed by affinity in the seed region, the lack of correlation between genomic localization and where the enzyme cleaves the genome suggest that this model is lacking.

Thus, despite significant progress in the identification of off-target cleavage and binding of the Cas9-RNP, the impact of mismatches on overall enzyme efficiency is not known. Modeling enzyme efficiency requires measurement of affinity and activity under the same conditions. To do this, we developed an *in vitro* approach to systematically measure both in parallel. We first used a technique we recently developed, *SELEX-seq* followed by generalized linear model fitting (*SelexGLM*) (Zhang, in revision, (23), to generate a biophysical model of Cas9-RNP binding specificity over a large (>30bp) binding site. To separately quantify the effect of mismatches on endonuclease activity and DNA-binding of the Cas9-RNP, we developed a new technique, Sequence-specific endonuclease activity measurement by sequencing (*SEAM-seq*), adapted Specificity measured by sequencing (*Spec-seq*) (24, 25), and developed new mechanistic models of Cas9-RNP specificity. Together, these approaches allowed us to identify regions where the effects are different or even opposing.

## MATERIAL AND METHODS

### Recombinant Cas9 expression and purification

The plasmid encoding WT-Cas9 (pMJ915) was a gift from Jennifer Doudna (Addgene plasmid # 69090). Point mutations (D10A and H840A) were created by site-directed mutagenesis to express catalytic deactivated Cas9 (dCas9) using primer pMJ915_D10A and pMJ915_H840A (See **Table S1** for sequence). Both Cas9 and dCas9 were expressed and purified as previously described (26). Briefly, the Cas9 protein was overexpressed in Rosetta 2 DE3 cells, and sequentially purified using Ni^2+^ affinity, cation exchange and size exclusion chromatography. N-terminal His- and MBP-tags were removed by TEV protease prior to the cation exchange chromatography. The purified protein was dialyzed into storage buffer (20 mM HEPES-KCl, pH7.5, 150 mM KCl, 10% Glycerol, and 1 mM TCEP), concentrated by Amicon ultrafiltration device (30-kDa), aliquoted, flash-frozen in liquid nitrogen, and stored at -80°C. The protein concentration was determined by Nanodrop using an extinction coefficient of 120,450 M^-1^cm^-1^.

### Cas9-RNP assembly

The NanogSg3 single guide RNA was synthesized (Synthego), and dissolved in the Cas9 storage buffer to a final concentration of 10 μM. The sgRNA was refolded by first incubating at 70°C for 5 minutes, and then gradually cold down to 25°C over 30 minutes. MgCl_2_ was added to a final concentration of 1 mM, followed by incubating at 50°C for 5 minutes, gradually cold down to 25°C over 30 minutes (27). Refolded sgRNA was aliquoted and stored at -80°C. To assemble the Cas9-RNP, 3 μM of purified Cas9 was incubated with 3.6 μM refolded sgRNA at 37°C for 10 minutes (20 mM HEPES-KOH, pH 7.5, 150 mM KCl, 10% Glycerol, 1 mM MgCl_2_, and 1 mM TCEP). Freshly assembled Cas9-RNP was used for each assay.

### SelexGLM

A DNA library with 30-bp randomized region (**Table S1)** was synthesized as single-stranded DNA by IDT using hand-mixing option. The double-stranded DNA library (dsDNA) was prepared by a Klenow extension reaction using Cy5-labeled TSSR1 primer (**Table S1**) that is complementary to the 3’-end of the ssDNA library. Briefly, a reaction containing 2.5 μM ssDNA template, 5 μM Cy5-labeled TSSR1, and 150 μM dNTP in NEB buffer 2 (10 mM TrisHCl, 50 mM NaCl, 10 mM MgCl_2_, 1 mM DTT) was incubated at 94°C for 3 minutes, and then slowly drop to 37°C over 45 minutes. Klenow enzyme was then added to the reaction, and incubated at 37°C for 1 hour. The enzyme was inactivated at 72°C for 20 minutes, followed by gradually cooling down to 10°C over 45 minutes. The dsDNA library was purified and concentrated (Qiagen MinElute). The concentration was measured by A260 on a Nanodrop, and then diluted to 4 μM in Qiagen buffer EB (10 mM Tris-HCl, pH 8.5).

*SELEX-seq* of dCas9-RNP was performed based on our previous protocol with slight modifications. Briefly, a 120 μl binding reaction containing 0.1 μM dCas9-RNP and 1 μM SELEX library (1:10 ratio) was incubated at 37°C for 1 hour (20 mM HEPES-KOH, pH7.5, 150 mM KCl, 10% Glycerol, 5 mM MgCl_2_, and 1 mM TCEP), then resolved on a 4∼20% gradient gel (1X TGM: 25 mM Tris-Base, 192 mM Glycine, 5 mM MgCl_2_, pH 8.3) at room temperature. The next library was then generated by amplifying the recovered DNA. The new library was then generated by purifying the regenerated library (Qiagen MinElute) then diluting to 4 μM for next round of *SELEX*. Five rounds of *SELEX* were performed in total. The initial and final libraries (R0 and R5) were deep-sequenced on an Illumina HiSeq at a read depth of ∼25-30 million reads per library.

The *SELEX-seq* data was processed by the *SelexGLM* and *SELEX* R packages as described previously (Zhang, in revision). Briefly, a Markov model of order 6 was constructed from the R0 probes using the selex.mm() function from the *SELEX* package, and an affinity table for k=18 was constructed using selex.affinities(). An initial position specific affinity matrix (PSAM) was constructed from the relative affinity of all single-base mutations of the optimal 18-mer (“NNNNAAGAWKGGAAGNGG”). The PSAM was then expanded to the desired size by adding 9 neutral columns on each side to estimate the specificity outside of PAM and protospacer, and used as a seed for *SelexGLM*. The subsequent iteration of *SelexGLM* contains two main steps. First, the current PSAM is used to compute the affinity of each 36-bp window on each DNA probe. If the highest affinity window is greater than 95% of the total affinity (sum of all windows), the probe is saved for generating new PSAM. Next, a logarithmic link function is used to fit the base identity of the optimal window on each probe against the probe counts normalized by the frequency in Round 0 library computed by a 6^th^-order Markov model. The regression coefficients are interpreted as the free energy differences ΔΔG, and used in the next round of iteration. This iterative process continues until the PSAM converges. The final model was plotted as motif logo using REDUCE Suite (28).

### Spec-seq

Each individual *Spec-seq* library was ordered as ssDNA from IDT, and pooled equally for Klenow extension as described previously. The dsDNA library was purified and concentrated (Qiagen MinElute PCR purification kit), and further size-selected on a 20% (19:1) 1X Tris-Glycine native gel. DNA band at correct size (74-bp) was excised, electroeluted, and purified. The concentration was quantified by A260 and diluted to 1 μM in buffer EB.

The *Spec-seq* of dCas9-RNP was performed essentially as a single round of *SELEX* experiment with slight modification. Briefly, a 40 μl binding reaction containing 200 nM dCas9-RNP and 50 nM *Spec-seq* DNA library (4:1 ratio) was incubated at 37°C for 1 hour, and resolved on a 4∼20% gradient gel. Both bound and unbound DNA were excised, electroeluted, purified, and diluted to 40 μl in buffer EB. Recovered DNA from each fraction was amplified using TSSR2 and TSSR-RPIX primers to prepare sequencing library.

### Sequence-specific Endonuclease Activity Measurement (SEAM-seq)

*SEAM-seq* of WT-Cas9-RNP (Cas9-RNP) was performed in a 50 μl reaction containing 50 nM *Spec-seq* DNA library with either 200 nM Cas9-RNP or reaction buffer as mock treated. After 1 hour of incubation at 37 °C, EDTA was added (60 mM final) to both digested and input samples, followed by addition of 10 μl Proteinase K (20 mg/ml, Thermo) to digest Cas9 for 30 minutes at room temperature. The remaining DNA was purified (Qiagen MinElute) and eluted in 40 μl. The full-length, uncut DNA library is PCR amplified and deep-sequenced.

## RESULTS

### The dCas9-RNP has substantial specificity over a 23bp footprint

To determine the DNA binding specificity of the catalytically deactivated dCas9-RNP over a large footprint in a comprehensive, unbiased fashion, we performed *SELEX-seq* (**Figure 1A**). Briefly, we first assembled dCas9 bound to a gRNA (dCas9-RNP) designed against the Nanog gene (12), a gRNA that has been shown to have thousands of off-target sites. We then incubated the dCas9-RNP (0.1 μM) with a 30-bp randomized library (1 μM), flanked by PCR primer sites. The bound fraction was separated on a native gel, recovered, amplified, and incubated with dCas9-RNP again in the same molar ratio. This isolation and enrichment was repeated for a total of five rounds after which the input and Round 5 libraries were sequenced. The clear enrichment of high-affinity sites from *SELEX-seq* Round 5 (R5, **Figure 1B)**, indicated that selection was successful.

**Figure 1:**
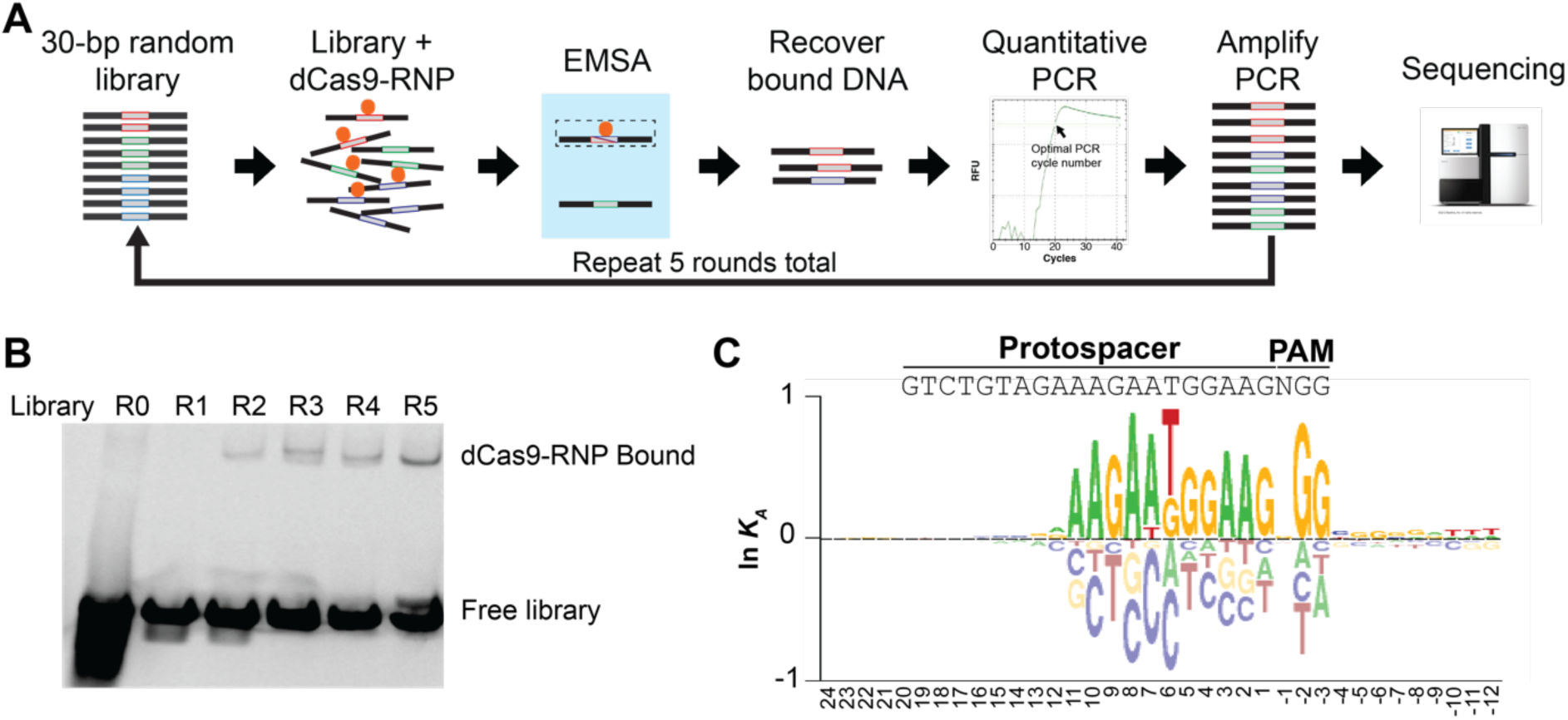
The DNA binding specificity of the dCas9-RNP extends over 23bp. **(A)** Schematic workflow of *SELEX-seq*. A 30bp random library flanked by sequencing adapters is incubated with the Cas9-RNP. Bound sequences are separated on an EMSA gel, excised, isolated, carefully amplified, and then used in the next round of selection. When performed under limiting Cas9-RNP concentrations, the round-by-round amplification is proportional to the affinity for a given sequence. **(B)** Summary EMSA gel. Equal concentrations of DNA library and Cas9-RNP are used in each lane. The increase in band intensity with each round indicates that the pool is being enriched for specific sequences. **(C)** Enrichment of sequences in the final library (R5) is compared to the initial library (R0) using a generalized linear model (*SelexGLM*) to derive a Position Specific Affinity Matrix (PSAM), reflecting the contribution of each base pair to the affinity of the Cas9-RNP for DNA. The majority of specificity is derived from the PAM (-1 to -3) and the PAM-proximal protospacer (1-11), although significant affinity is derived from PAM-distal base paring (positions 12,13) and sequence downstream of the PAM (-4 to -12).

We processed the *SELEX-seq* data using the *SelexGLM* R package (Zhang, in revision). This software package uses a biophysical model of protein binding to infer how mononucleotide features within the binding site contribute to the free energy of binding. The resulting position-specific affinity matrix (PSAM) encodes the energetic contribution of each nucleotide to affinity, and can be visualized as an energy logo (29). From this PSAM, the relative *K*_*D*_ (or *K*_*A*_) can be calculated for any sequence by adding the differences in free energy from mismatches compared to the perfect match. *SelexGLM* can model the binding affinity over a large footprint, which is essential for understanding dCas9-RNP binding.

The energy logo for the dCas9-RNP has two striking features (**Figure 1C**): 1) the expected preference of the dCas9-RNP for an NGG PAM site; and 2) the lack of sequence preference in the PAM distal region (13-20) of the protospacer. There is a clearly defined region of sequence preference within the PAM-proximal protospacer that is consistent with a seed sequence confined to the PAM-proximal driving binding. Most specificity is contained in PAM-proximal bases 1-11, with mild preference for matches at positions 12-13. Within the PAM-proximal region, the greatest binding energy is derived from exact matches, with some tolerance at positions 6 and 7 for a G and T mismatch, respectively. The dCas9-RNP also exhibits some preference for Gs downstream of the PAM (–5 to –8) and Ts further downstream (–10 to –12). Although minor compared to the affinity contributed by the PAM and PAM-proximal base pairs (∼6% *vs.* 92% of total specificity computed by maximal ΔΔG/RT), these downstream sequences contribute more to DNA binding affinity than PAM-distal base pairs (0.5 kcal/mol vs. 0.2 kcal/mol). Despite the lack of preference in the PAM-distal region, the dCas9-RNP nonetheless is sensitive to base identity over a 23-bp footprint.

The PSAM (**Figure 1C**) not only reveals which bases are preferred by the dCas9-RNP, but also the mismatches that are most disfavored. The gRNA used in this experiment is highly purine-rich within the protospacer (15/20 positions, including 12/3 within the PAM-proximal region). At these purine positions, the most detrimental mismatches are non-complementary pyrimidines (A->C, G->T). These mismatches, for example C instead of A, would result in the A of the gRNA attempting to base pair with a G on the opposite strand, which is both non-complementary and bulky. This pattern is remarkably consistent, and indicates that dG·rA and dA·rG base pairing between the gRNA and protospacer is the most detrimental to binding. An exception to this is the T at position 6, which also tolerates a C least. This C would also result in a non-complementary G opposite the T in the gRNA. Together, it is clear from these data that not all mismatches have an equal effect on binding, with some more detrimental than others.

### Parallel measurement of DNA binding and cleavage using Spec-seq and SEAM-seq

Cas9-RNP activity involves the two-step process of binding followed by cutting DNA (**Figure 2A)**. The overall efficiency of the enzyme depends on how well it binds a sequence (*K*_*A*_) then how quickly it is cut (turnover or *k*_*cat*_). To generate these two values, we paired the techniques of *Spec-seq* (24, 25) to measure affinity and *SEAM-seq* (this work) to measure endonuclease activity (**Figure 2B,C**, described in detail below). Performing these assays under identical conditions on a single library, composed of seven sub-libraries encompassing a total of 1,594 sequences, allowed direct comparison of the effect of mismatches on binding and nuclease activity for each sequence. By fitting both data sets to sequence features, specificity models for both binding and cleavage can be generated. Further, we are able to estimate the relative occupancy of sequences based on the measured affinity in *Spec-seq*, calculate the turnover of the enzyme, and generate a general model for the enzyme efficiency throughout the protospacer, PAM, and flanking regions. This general efficiency model can then be used to estimate off-target cutting at sites that differ from the intended target.

**Figure 2.**
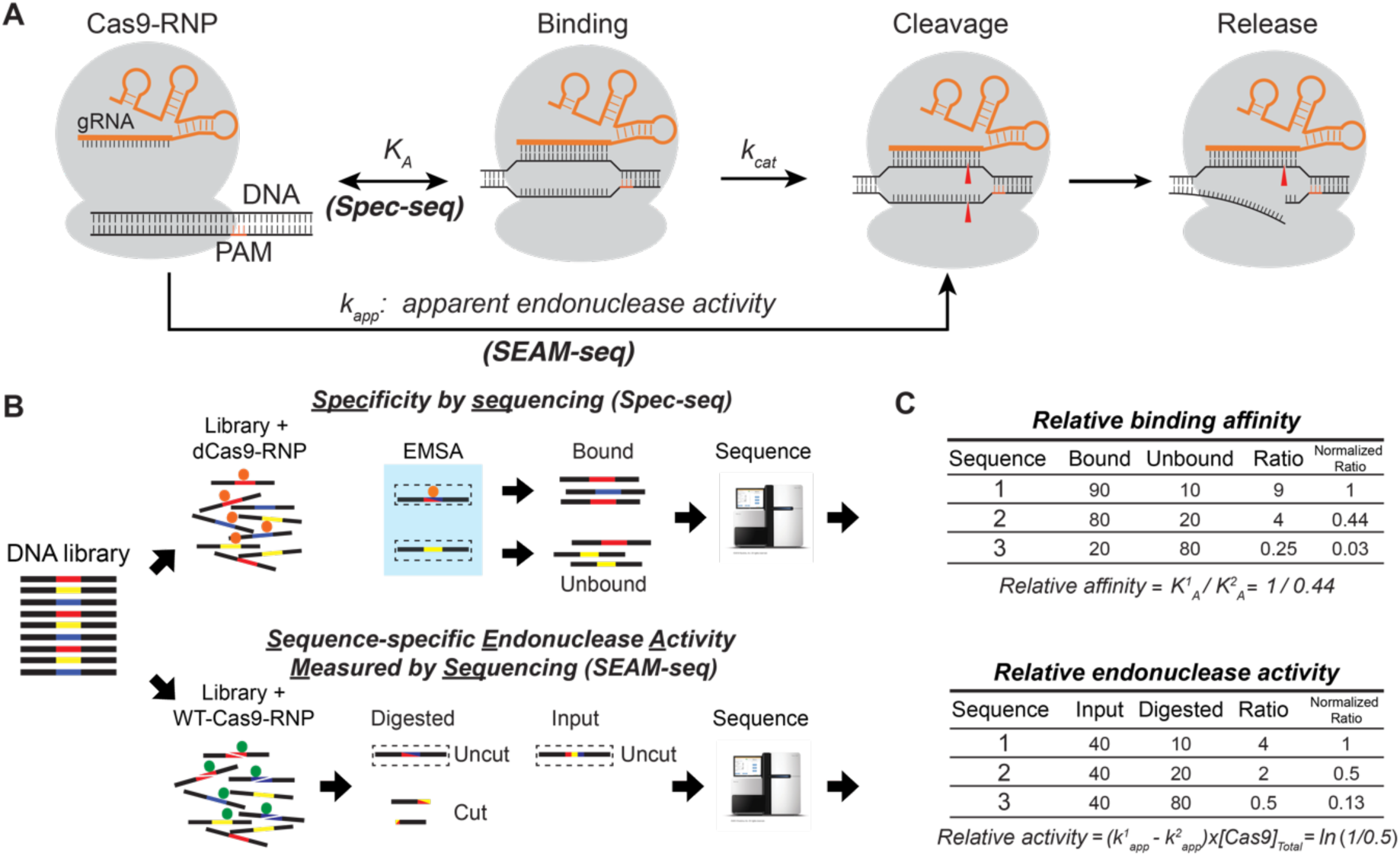
Decoupling DNA binding and endonuclease activity of the Cas9-RNP by *Spec-seq* and *SEAM-seq*. **(A)** Two-step enzymatic model of Cas9-RNP-mediated DNA cleavage. The apparent endonuclease activity of the Cas9-RNP (*k*_*app*_) is determined using *SEAM-se*q, whereas the relative affinity of binding (*K*_*a*_) is measured by *Spec-seq* for each of the 1,594 sequences in the library. **(B,C)** Schematic of the *Spec-seq* and *SEAM-seq* protocols. For *Spec-seq*, sequences bound by the catalytic-deactivated Cas9-RNP are resolved from unbound on an EMSA gel. Each band is excised and deep-sequenced, with the relative affinity directly calculated **(C)** from the ratio of bound to unbound (see Materials and Methods). For *SEAM-seq*, the same sequences are cleaved by wild-type Cas9-RNP. Uncut sequences are amplified by PCR and deep-sequenced. The ratio of uncut sequences in digested to input fraction reflects the relative cleavage of sequences **(C)**, which can be normalized to calculate the relative apparent endonuclease activity (Δ*k*_*app*_, see Materials and Methods).

*The SELEX-seq and Spec-seq binding specificity measurements are consistent.* To measure the relative affinity of the dCas9-RNP for each sequence, we performed standard *Spec-seq* (24, 25) using the dCas9-RNP. Sub-libraries were pooled equally and incubated with dCas9-RNP at a 1:4 ratio (50 nM DNA: 200 nM RNP). Bound and unbound fractions were then separated on a native gel (**Figure S1A**), recovered, and sequenced. After ensuring that sub-libraries are similarly represented in the full library (**Figure S2C**), the relative binding affinity of dCas9-RNP for each site were calculated directly by taking the ratio of the probe frequencies in the bound and unbound fractions (See Methods). At a sequencing depth of ∼30 million reads per sample, each sequence was represented by an average of ∼20,000 counts. This allowed quantification of affinities across four orders of magnitude, as compared to ∼2 orders of magnitude for *SELEX-seq* followed by *SelexGLM* analysis (**Figure 3C, E, Figure S1C**).

**Figure 3.**
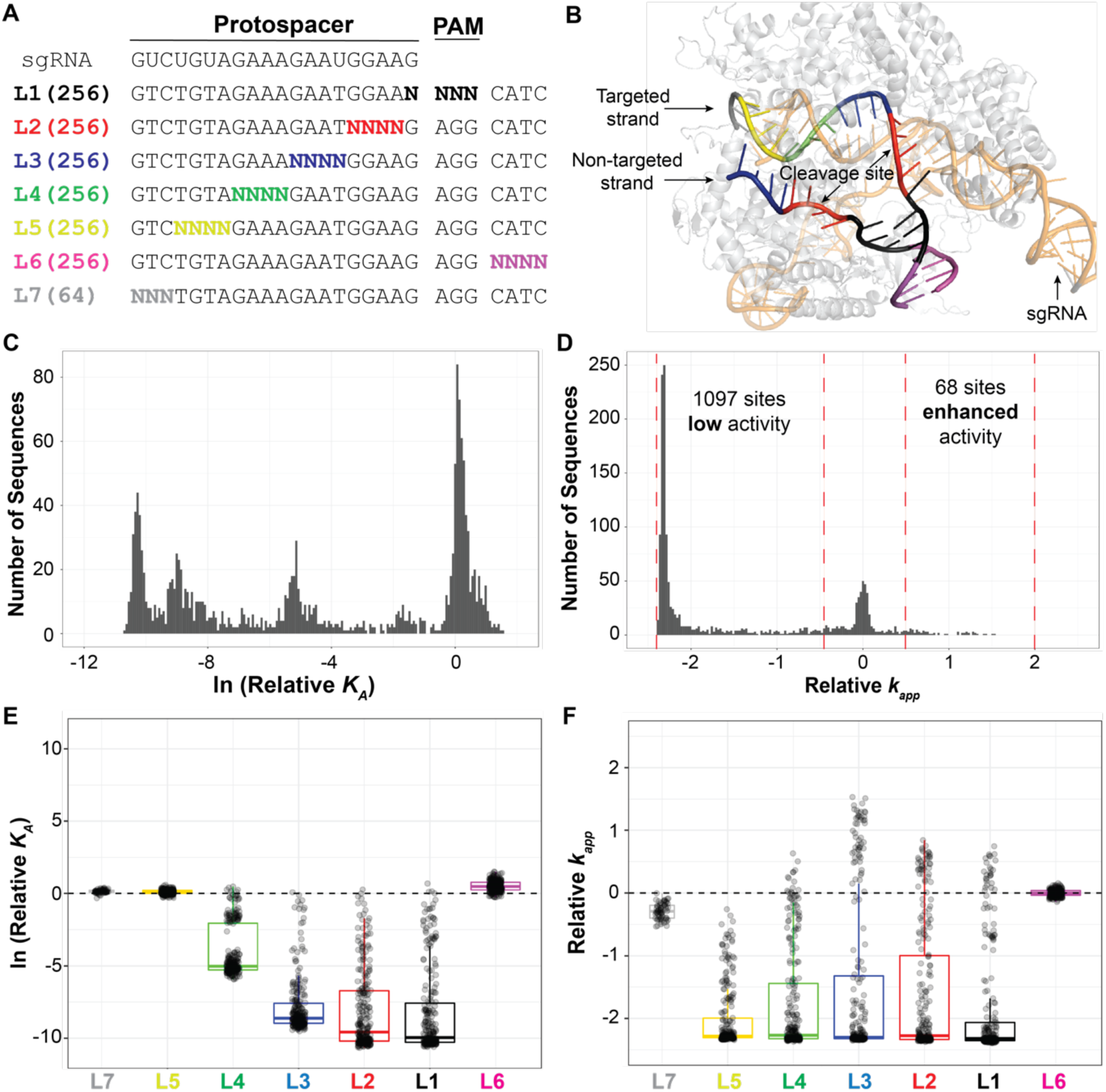
The sequence preference for endonuclease activity and affinity is similar, except in the PAM-distal region. **(A)** Seven sub-libraries were pooled to comprise the library used in *SEAM-seq* and *Spec-seq*. The sequences cover all possible single mismatches and blocks of up to 4 mismatches through and downstream of the PAM as well as the protospacer. **(B)** DNA libraries mapped on the existing crystal structure of Cas9-RNP:DNA complex (PDB: 5F9R). L3 is positioned at the bend in RNA:DNA duplex. **(C)** Histogram showing the distribution of relative binding affinity indicating that many mismatches to not cause a large effect on binding. **(D)** Histogram of the relative apparent endonuclease activity indicating that most mismatches decrease activity, although some do increase cutting (farther right). **(E, F)** Boxplots showing the distribution of the effect of mismatches on affinity and endonuclease activity. **(E)** Mismatches closer to the PAM have a more deleterious effect than distal mismatches on affinity. **(F)** Many mismatches increase endonuclease activity, particularly in L3 (positions 6-9).

The results obtained using *Spec-seq* agreed very well with the PSAM obtained using *SelexGLM*. In particular, we observed a high correlation between the directly measured *Spec-seq* sequences compared to the calculated affinity from the *SelexGLM* PSAM (Spearman ρ = 0.88, R^2^ = 0.85, **Figure S1C**). Inconsistencies between the two methods are likely due to two main factors: sequencing depth and any non-additive effect of multiple mismatches on affinity. Capturing the ∼10,000-fold difference in affinity measured by *Spec-seq* requires differences in reads for *SELEX-seq* scaled to the round of selection (R5 in this case), or a sequencing depth of 10^20^. As a result, low-affinity sequences are not captured by *SelexGLM*, resulting in apparent compression compared to *Spec-seq*. Also, because *SelexGLM* assumes that all positions are independent, it likely underestimates the effect of multiple mismatches, whereas these effects are directly measured by *Spec-seq*. Despite these differences, a linear regression model fit to the *Spec-seq* data **(Figure S4A, S6B)** and the *SelexGLM* model **(Figure 1C, S6A)** are remarkably consistent. Thus, like the *SelexGLM* model built from *SELEX-seq* data, *Spec-seq* analysis confirms that purine:purine mismatches are the least favorable for binding, the PAM-distal region does not contribute to affinity, and that the closer a mismatch is to the PAM, the more deleterious the effect is on binding (**Figure 3E**). However, there are also exceptions. For example, some PAM-proximal mismatches have very little effect on affinity (Libraries 1, 2, and 3, **Figure 3E**), such as the change of guanine at position of 4 of protospacer to adenine (G4A), T6A, and G9A (ln*K*_*A*_: -0.01±0.27, 0.08±0.26, -0.12± 0.06 respectively). These examples are reproducible, indicating that some PAM-proximal region mismatches can be tolerated by the dCas9-RNP.

### Sequence-specific endonuclease activity measured by sequencing (SEAM-seq)

To accurately measure the relative endonuclease activity of WT-Cas9-RNP (Cas9-RNP), we developed *SEAM-seq*, a method explicitly designed to be paired with *Spec-seq* (24, 25) (**Figure 2B**). First, the same *Spec-seq* library (50 nM) was used to measure the binding specificity of the dCas9-RNP is incubated with a 4x excess of Cas9-RNP (200 nM), and allowed to cleave for 1 hour under the same conditions that we ran *Spec-seq*. At this nuclease:DNA ratio, high affinity sites (K_D_∼1 nM to 100 nM) are expected to be fully occupied by the enzyme, whereas lower affinity sites (K_D_ > ∼100 nM) only partially occupied. At the end of incubation, the remaining uncut DNA was amplified by PCR. A control mock digestion (input) was performed by incubating library with buffer, and subjected to the same downstream processes. Both the uncut and the input libraries were then sequenced. With each library being similarly represented (**Figure S2**), the ratio of reads between uncut and input libraries measures the relative fraction cut, or relative depletion of each DNA sequence by the Cas9-RNP. The logarithm of this depletion ratio is proportional to the difference of apparent relative rate of cleavage (or Δ*k*_*app*_) (**Figure 2B, C;** see Method section for details). To establish a reference baseline for the relative cut rate, we set the average cut rate of all exact matches within the PAM and protospacer to zero, and calculated the relative impact of mismatches on the endonuclease activity. Measured in this way, sequences with a *k*_*app*_ < 0 are cleaved less than the perfect match, whereas those with a *k*_*app*_ > 0 are cleaved more during the 1 hour incubation. Two experimental replicates of *SEAM-seq* showed that the technique is highly reproducible (**Figure S1E**, Spearman’s ρ = 0.93, R^2^ = 0.96), with the average *k*_*app*_ used for downstream analyses.

### Base pairing throughout the protospacer is important for endonuclease activity

To investigate how mismatches disrupt endonuclease activity, we modeled the endonuclease activity using linear regression with mononucleotides features as predictors (**Figure S5A**). This revealed that endonuclease activity requires base pairing over almost the entire protospacer (mismatches at position 20 have a less prominent effect) (**Figure 3D, F, S5A**). This is consistent with the previous model where base pairing in the PAM-distal region is critical for endonuclease activity (7), and in marked contrast to the more localized effect of each nucleotide on binding affinity (1). Analogous to their effect on binding affinity, the sequence logo revealed that purine:purine mismatches tend to disrupt endonuclease activity most strongly. Lastly, although sequences downstream of the PAM contribute to affinity, variation in this region does not affect endonuclease activity significantly.

### Most combinations of mismatches in the PAM-proximal region decrease affinity and activity

Plotting the binding affinity measured by *Spec-seq* against the endonuclease activity measured by *SEAM-seq* showed that most (871 out of 1007) sequences with reduced binding also exhibited reduced nuclease activity (**Figure 4A**). This correlation is, as expected, strongest for mismatches within the PAM and the PAM-proximal region (positions -3 to +11, black, red, blue, and part of green libraries) where mismatches result in lower affinity (**Figure 4B-E**). This correlation between affinity and activity is most striking when the affinity is ∼150x less (ln*K*_*A*_ < -5) than the perfect complement. At the concentration used in the experiment, and the K_D_ of the Cas9-RNPs for the perfect complement is ∼1 nM, the reduced endonuclease activity over those low affinity sites is primarily due to reduced occupancy.

**Figure 4:**
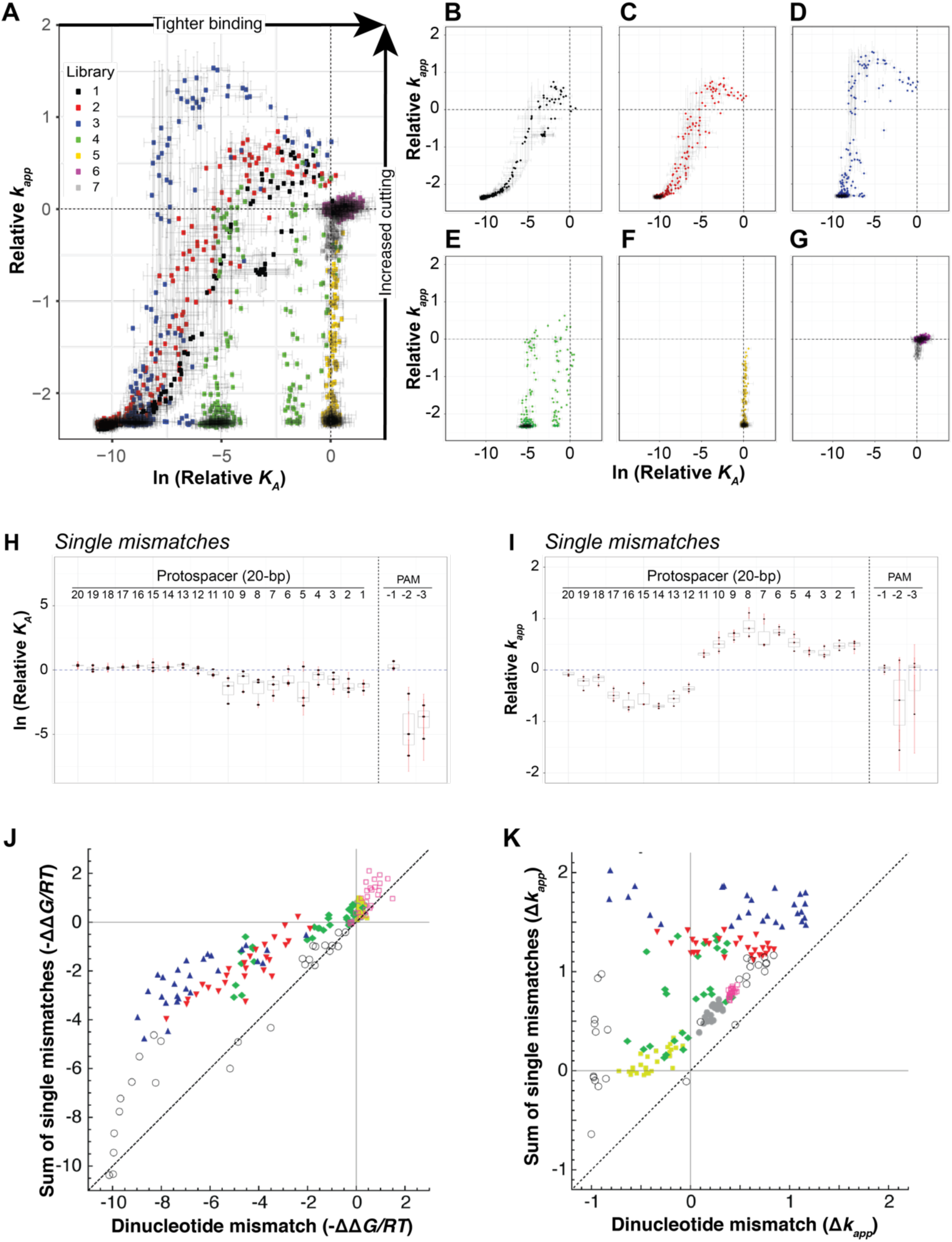
Dozens of sequences increase the endonuclease activity of the Cas9-RNP. **(A)** Scatterplots of the relative binding affinity (x-axis) *versus* the relative endonuclease activity (y-axis) for all sequences, and **(B-G)** for each library separately. In general, mismatches decrease the affinity and the endonuclease activity toward the lower left. However, mismatches in the PAM-proximal region can often increase the endonuclease activity (y-axis > 0) while decreasing the affinity (particularly apparent in **B-E**). Error bars represent the standard deviation based on two *Spec-seq* and *SEAM-seq* repeats. **(H,I)** Plot of the effect of single mismatches on binding affinity and endonuclease activity. The penalty of single mismatch on affinity is relatively low (< 20-fold), even in the PAM-proximal region. However, **(I)** all single mismatches in the PAM-proximal region (1-11) increased endonuclease activity whereas all mismatches in PAM-distal region (12-20) reduced the endonuclease activity.

### Affinity does not always correlate with endonuclease activity

Three classes of mismatches do not have corresponding reductions in affinity and activity: 1) those at PAM-distal positions (+12 to +20); 2) those downstream of the PAM (-4 to -8); and 3) single and multiple mismatches in the PAM-proximal region that enhance endonuclease activity. First, consistent with previous observations (7), mismatches in the PAM-distal region (12-20, yellow, grey and part of green libraries) have little effect on binding affinity, but universally impair endonuclease activity (**Figure 3F**). Second, specific sequences downstream of the PAM enhance affinity but do not affect activity. These sequences, if present in the genome, would have enhanced endonuclease activity, but can be easily identified and avoided.

Most interesting is the third class, which consist of mismatches in the PAM-proximal region that enhance endonuclease activity, a phenomenon we term mismatch-activation. These sequences can be most easily visualized in the breakout plots for each individual library (black, red, blue, part of green, positions -3 to 11, and **Figure 4B-G**). As discussed above, most mismatches within this region (649 out of 781) impair both binding and endonuclease activity and fall along a binding sigmoid from the origin to the lower left of the plots. However, several sequences lie above this sigmoid, indicating enhanced endonuclease activity. Further, some of these sequences are cut more completely than even the perfect match (*k*_*app*_ > 0), even while affinity is lower (relative ln*K*_*A*_ < 0). A group of these mismatch-activated sequences is particularly evident in the blue library (positions 6 to 9). Within this library, the activity against some sequences are ∼1.5x greater than the perfect match, indicating that sequences that don’t match the gRNA can strongly enhance enzyme activity (**Figure S2A**). Among these mismatch-activated sequences, 26 of 27 have one or two mismatches, with the remaining sequence having three mismatches (**Figure S2B**). Under the conditions of the experiment, the mismatch-activation sequences are likely to be fully occupied by the Cas9-RNP, indicating that the increased activity is due to increased turnover and not affinity.

Single mismatches in the PAM proximal region almost universally enhance Cas9-RNP endonuclease activity (**Figure 4I**). This single base pair mismatch-activation reaches a peak with mismatches between positions 3 and 9 (**Figure 4I**), but remains significant at other positions in the region. Though activating, these single mismatches nonetheless reduce affinity (**Figure 4H**). By contrast, pairs of adjacent mismatches throughout this region reduce affinity synergistically (**Figure 4J**), whereas the same pairs have a less that additive effect on mismatch-activation (**Figure 4K**). These findings suggest that destabilizing the RNA:DNA helix can enhance enzymatic activity even while reducing affinity. When the helix is destabilized further by adjacent mismatches, enhancement is still observed, but the penalty for binding continues to increase, reducing the probability of off-target cutting.

### The addition of DNA features improves the Spec-seq affinity model

Although *Spec-seq* and *SEAM-seq* directly measure the activity of ∼1,600 sequences, few of these are found in the genome. We therefore sought to develop general models that could predict off-target binding and cutting for sequences not present in our library. To predict the binding affinity, we first assumed that the binding free energy ΔΔG is due to independent linear contributions from each nucleotide in the binding site (described above). This generated a PSAM (**Figure 5A**) that agreed well with *SELEX-seq* PSAM and the model predicted the measured ΔΔG values quite well (**Figure S4A**, model *Mono*, *R*^2^ = 0.95). However, plotting predicted *versus* measured ΔΔG values revealed significant model bias; the affinity of low-affinity probes was underestimated and the range of predicted valued for higher-affinity probes was compressed (**Figure S4A**). We attempted to improve our model by adding a non-specific binding component that moderates the predicted affinity of poorly matched sequences. This dramatically improved the fit and removed most of the bias (**Figure S4B**, model *Mono/NS*, *R*^2^ = 0.98). As noted above, we observed a non-linear contribution of adjacent dinucleotide mismatches to binding (**Figure 4J**). When we incorporated the dinucleotide interactions into the model, the fit improved still further (**Figure 5A, S4C**, model *M/Di/NS*, R^2^ = 0.99). This method not only generates a very accurate model for the effect of any mismatch on binding but also validates the finding that adjacent mismatches have a greater than additive effect on affinity (8).

**Figure 5.**
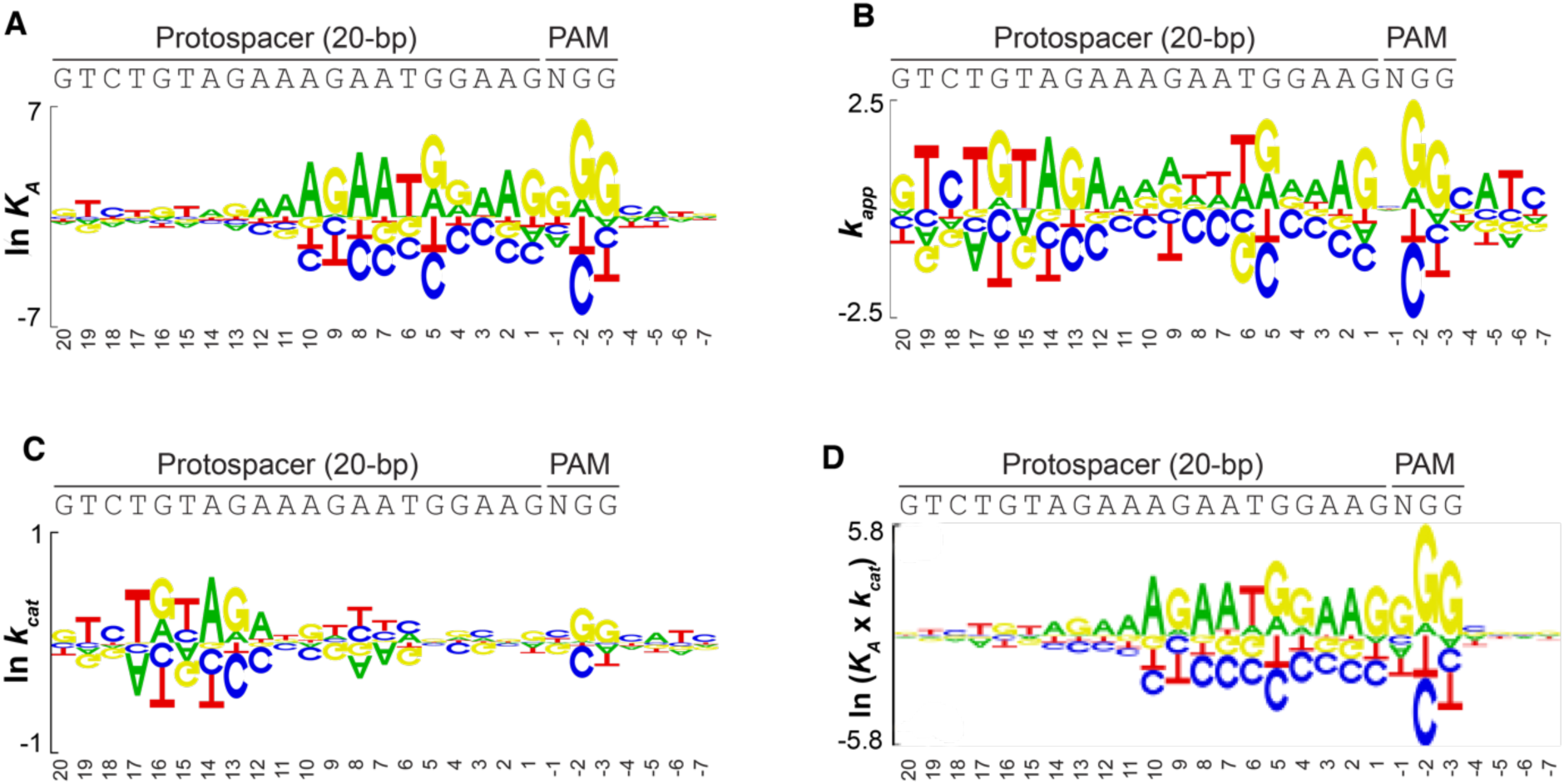
Accurate modeling both binding affinity (*K*_*A*_) and apparent endonuclease activity (*k*_*app*_). Sequence logos showing how the different bases contribute to the binding affinity **(A)** and apparent endonuclease activity **(B)** using regression models that incorporate the contribution of mononucleotide and dinucleotide features, and corrected for non-specific binding. **(C)** Sequence logo depicting the sequence dependence of endonuclease activity corrected for occupancy activity (*k*_*cat*_) derived using both *SEAM-seq* and *Spec-seq* data. **(D)** The enzymatic efficiency model of Cas9, which is the product of *K*_*A*_ and *k*_*cat*_.

**Figure 6.**
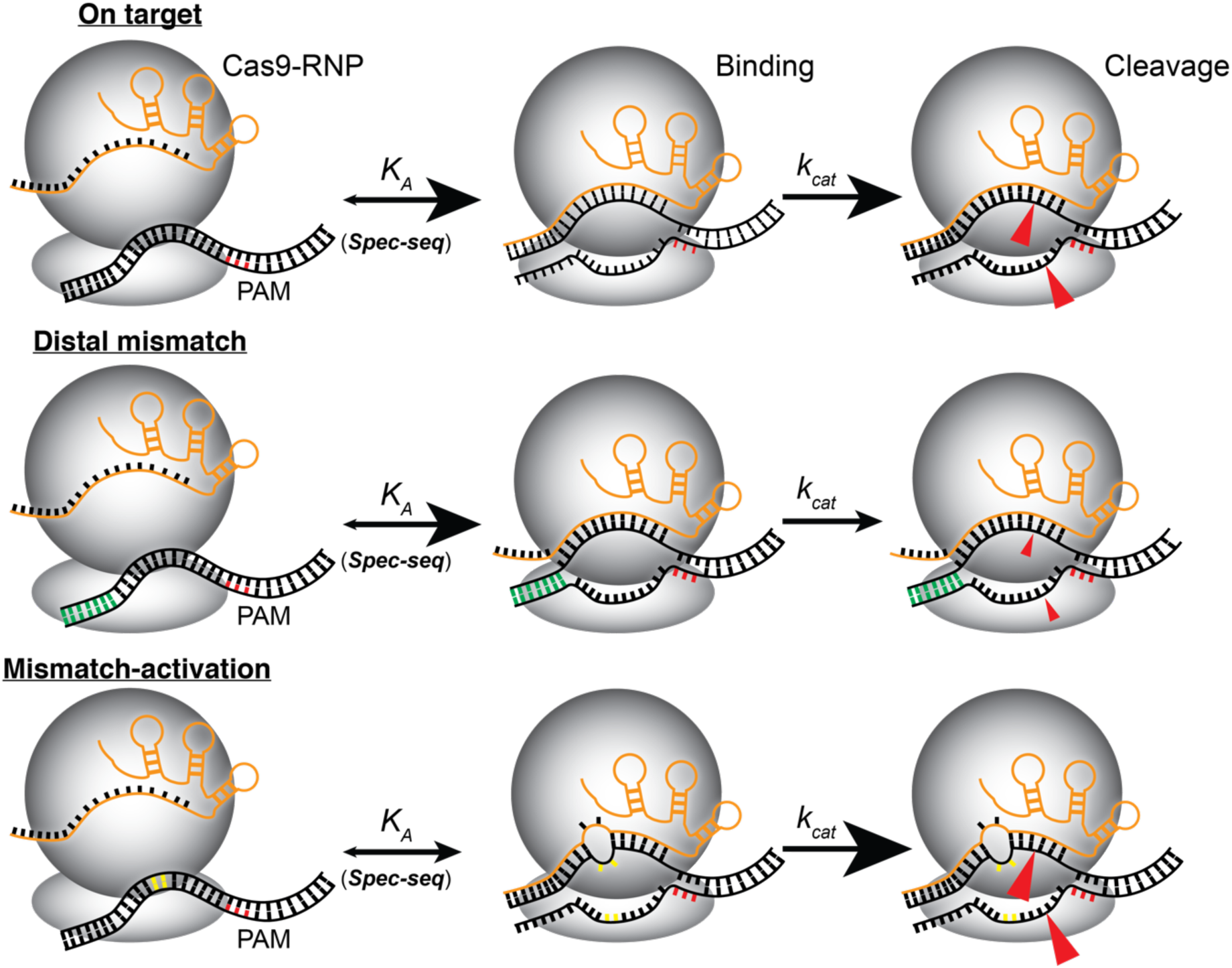
Distinct impact of mismatches on DNA binding and cleavage activity of the Cas9-RNP. Models summarizing the effect of sequence on both affinity (*K*_*A*_) and turnover (*k*_*cat*_) of the enzyme. The perfect match (top) has both high affinity and activity. Mismatches in the PAM-distal region (green) do not affect affinity, but severely impair *k*_*cat*_. Single and double mismatches within the PAM-proximal region (yellow) can decrease affinity, but substantially increase turnover (*k*_*cat*_).

### Calculating the catalytic activity of Cas9 by integrating Spec-seq and SEAM-seq

To generate a general model for apparent endonuclease activity (*k*_*app*_) we started with a linear model that used only mononucleotide features. This simple model described the *SEAM-seq* data reasonably well but also exhibited significant bias (**Figure S5A**, model *Mono*, *R*^2^ = 0.77), suggesting that a more refined model of *k*_*app*_ was needed. The *k*_*app*_ measured by *SEAM-seq* reflects a two-step process (**Figure 2A**): binding (*K*_*A*_) followed by affinity-independent cutting (*k*_*cat*_). Because the binding of higher affinity sequences was more saturated than lower affinity sequences, we reasoned that correcting for occupancy would enable estimation of *k*_*cat*_. Having run the *SEAM-seq* and *Spec-seq* experiments under matched experimental conditions, we were able to determine the relative occupancy of Cas9 by assuming that the occupancy has a sigmoidal dependence on the binding affinity and included the free protein concentration as a free parameter. This allowed calculation of model for *k*_*cat*_, and allowed generation of a model for Cas9-RNP efficiency for off-target sequences.

The model for *k*_*cat*_ (**Figure 5C**), which represents affinity-independent cleavage, significantly improved our modeling of observed *k*_*app*_, when used in combination with the relative occupancy predicted by affinity model (**Figure S5E, bottom plot**). As expected, because mismatches do not affect affinity in the PAM-distal region, the sequence logo of *k*_*cat*_ shows a negative effect on activity for any mismatches in the PAM-distal region. Conversely, purine:purine mismatches from positions 6-9 within the PAM-proximal region have increased activity, mirroring the mismatch-activation described above. Surprisingly, the mismatches with the greatest activation (C and T) exhibit the most negative effect on affinity. Thus, although mismatches can enhance activity of the Cas9-RNP, increased off-target cleavage is offset by decreased binding to some extent. Finally, the improved performance of the model suggests that it can be used to predict the *k*_*app*_ for the Cas9-RNP at different protein concentrations.

## DISCUSSION

As Cas9 and other RNA programmed endonucleases are increasingly widely used in genomic experiments, there is a pressing need to be able to predict off-target activity. The specificity of an enzyme is a function of the efficiency for off-target sites and the concentration of enzyme and ligand. Thus, predicting off-target cutting requires a measurement of efficiency, or the product for the turnover rate of the enzyme (*k*_*cat*_) and the propensity to bind (estimated by *K*_*A*_). Previous experiments have identified sites that can be cut in the genome or from random libraries (17, 19, 30, 31), and have also identified alternate binding sites from genomic experiments (12, 13). Although these have provided information about where the Cas9-RNP can cut, no clear rule can be summarized from any of these assays to predict where the Cas9-RNP will cut. Here we present a systematic experimental and computational pipeline to measure Cas9-RNP efficiency for on- and off-target sites that can be performed rapidly for any gRNA.

To accurately measure the specificity of a Cas9-RNP, we produced the first free-energy based DNA binding model. Using a gRNA directed at the Nanog gene, which is known to have thousands off-target binding sites (12), we used our *SELEX-seq/SelexGLM* pipeline (Zhang, in revision, (23) to calculate a comprehensive, unbiased PSAM model for binding any sequence. This model, from which the relative *K*_*A*_ of any sequence can be calculated, confirmed and provided more details than previous models, reinforcing the idea that the PAM, and PAM-proximal sequences (positions 1-13) are the determinants of binding affinity. The model also revealed a region downstream of the PAM that contributes to specificity in sequence-dependent fashion. Finally, the PSAM also revealed, as had been suggested previously (16, 32), that not all mismatches have the same effect on affinity, with purine:purine mismatches between gRNA and targeted strand of DNA being the most deleterious. Importantly, this model was recapitulated using a much simpler and less labor-intensive technique, *Spec-seq*, that can be paired with our endonuclease activity assay, *SEAM-seq*.

In addition to thoroughly validating the *SelexGLM* PSAM model, the *Spec-seq* results provided insight into the mechanism of binding. Due to the depth of sequencing, we were able to use linear regression to incorporate the contribution of non-specific binding and dinucleotide pairs to binding, which revealed that the thermodynamic contribution of each base pair to binding is not independent (**Figure 4J, 5C**). This finding and the diminished contribution of complementarity to affinity as distance from the PAM increases from positions 10-13 support the model that PAM-melting is followed by directional sequential base-pairing over ∼13 sites to achieve full affinity (8). Overall, these models are a significant improvement over previous *ChIP-seq* based methods (**Figure S6C**), and can be used to estimate binding at off-target sites, even those that contain mutations (12). More importantly, the binding specificity of the dCas9-RNP suggests that a re-evaluation of how gRNAs are designed for dCas9-based genomic experiments is needed.

Binding to DNA is only the first of two steps required for DNA editing by Cas9-RNP. By pairing the relative *K*_*A*_ values with apparent endonuclease activity (*k*_*app*_) using *SEAM-seq*, it is now possible to decouple the effects of mismatches on affinity and activity. Although in most cases the two steps are correlated (e.g. decreased affinity leads to decreased cutting), there are systematic exceptions that shed light on the mechanism. First, as has been observed previously, it is clear that mismatches in the PAM-distal region impair enzymatic activity while affinity is unaffected. Interestingly, multiple mismatches have a less than additive effect on activity, indicating that an initial break in base pairing is the key step (**Figure 4K**). Second, and perhaps most strikingly, mismatches within the PAM-proximal region of protospacer, while they decrease affinity, actually enhance enzymatic activity. Importantly, adjacent mismatches within this region decrease binding synergistically, while enhancing activity less than additively. This indicates that the loss in binding affinity of mismatches can be partially compensated by enhanced activity. This has implications in designing gRNAs to cut specifically in the genome, but also reveals that the Cas9 system for bacterial defense against viral invasion has a built-in tolerance for mismatches in the PAM-proximal region of the protospacer, possibly due to an evolutionary pressure to adapt to mutations in viral genomes. These findings, that binding does not always result in efficient cutting, help to explain the lack of overlap between where the Cas9-RNP binds the genome, and where it cuts (12, 13, 30).

Mismatch activation is particularly prominent between positions 6 and 9 of the PAM-proximal region. Structural studies have shown that the DNA:RNA duplex takes a sharp bend in this region (**Figure 3B**). Mismatch-activation has been noted in previous experiments using other two different gRNAs (33). Although not observed as consistently, this phenomenon was nonetheless most prominent in same region (positions ∼5-10) (33). However, it was not known at that time whether the increased cutting was due to increased enzyme turnover or enhanced binding. Whether mismatches enhance activity by facilitating this bend awaits further study.

Finally, with a pipeline to measure affinity and activity under the same conditions in hand, the relative efficiency of a Cas9-RNP can now be estimated using regression-based modeling of *Spec-seq* and *SEAM-seq* data. By correcting for baseline activity and occupancy, *k*_*cat*_ can be calculated, enzymatic efficiency can be calculated as the product of *K*_*A*_ and *k*_*cat*_. Based on this model, it is clear that mismatches in the PAM-distal region create an enzyme that is less efficient. It is also clear that, although mismatches within the PAM-proximal region enhance activity, they cause similar overall decreases in efficiency. As a result, any single mismatch in the PAM-distal and late (positions 6-9) PAM-proximal region can have similar effects on efficiency. More PAM-proximal mismatches are even less efficient. Thus, in designing gRNAs, those with other genomic sites containing mismatches in the near PAM-proximal region will have a lower probability for off-target cutting than those in the more PAM-distal regions, which still retain significant efficiency.

In addition to providing insight into how mismatches affect the mechanism of Cas9-RNP binding and cleavage, we describe a simple experimental protocol and pipeline for determining the specificity of Cas9:gRNA pair. Because the *SELEX-seq* generated PSAM is recapitulated so well by the *Spec-seq* model for binding, it is reasonable to design libraries similar to those described here based on gRNA sequence alone. Compared to other high-throughput measurements of binding (such as: CHAMP(34) or Hits-Flip(35)), *Spec-seq* and *SEAM-seq* require no special equipment, and multiple gRNAs can be assessed in parallel as described with sequencing-ready libraries in a few days. The computational demands are also minimal, and the scripts provided can be run on the resulting data, with results in a few hours. The robust models generated by this technique will allow estimation of off-target binding and cleavage for any gRNA, modified Cas9, or other sequence-specific endonuclease to guide the design of the most appropriate tool for specific applications.

## ACCESSION NUMBERS

Sequencing reads will be deposited prior to publication.

## ACKNOWLEDGEMENT

We thank members of the Pufall and Bussemaker labs for useful discussions. We also thank Dipa Sashital and Stephen Floor for critically reading the manuscript and providing helpful input.

## FUNDING

This research was supported by the National Science Foundation CAREER grant 1552862 (M.A.P), and grant R01HG003008 (H.J.B) from National Institutes of Health.

## CONFLICT OF INTEREST

The authors have no competing financial of personal interests to declare.

## SUPPLEMENTARY DATA

Supplementary figures and data can be found in the attached Supplementary document.

